# The Soft Vertex Classification for Active Module Identification Problem

**DOI:** 10.1101/407460

**Authors:** Nikita Alexeev, Javlon Isomurodov, Gennady Korotkevich, Alexey Sergushichev

## Abstract

**Motivation:** Integrative network methods are commonly used for interpretation of high-throughput experimental biological data: transcriptomics, proteomics, metabolomics and others. One of the common approaches consists in finding a connected subnetwork of a global interaction network that best encompasses significant individual changes in the data and represents a so-called active module. Usually methods implementing this approach find a single subnetwork and thus solve a hard classification problem for vertices. This subnetwork inherently contains erroneous vertices, while no instrument is provided to estimate the confidence level of any particular vertex inclusion. To address this issue, in the current study we consider the active module problem as a soft classification problem. We propose a method to estimate probabilities of each vertex to belong to the active module based on Markov chain Monte Carlo subnetwork sampling.

**Results:** The proposed method allows to estimate the probability that an individual vertex belongs to the active module as well as the false discovery rate (FDR) for a given set of vertices. Given the estimated probabilities, it becomes possible to provide a connected subgraph in a consistent manner for any given FDR level: no vertex can disappear when the FDR level is relaxed. We show on simulated dataset that the proposed method has good computational performance and high classification accuracy. As an example of the performance of our method on real data, we run it on a protein-protein interaction network together with a gene expression DLBCL dataset. The results are consistent with the previous studies while, at the same time, the proposed approach is more flexible. Source code is available at **https://github.com/ctlab/mcmcRanking** under MIT licence.

## 1. Introduction

Integrative network approaches are commonly used for interpretation of high-throughput data [13]. Such methods are applied in many different contexts: in genome-wide association studies [15], for elucidating mechanisms of metabolic regulation [10], for analysis of somatic mutations in cancer [12], etc. The main idea of these methods is that considering internal connections (for example, between proteins, metabolites or other entities) can lead to deeper understanding of the data and the corresponding biological processes.

The connectivity information could be used in multiple ways. The simplest analysis could involve manual exploring of connections between the input signals [11]. More sophisticated methods include using connections for gene set enrichment analysis [1], comparing networks [7] and many others.

One of the most well-developed and used approaches consists in selecting a connected subnetwork that best represents an *active* or *functional* module. This concept was initially suggested by Ideker et al [8]. The authors proposed a metric to score subnetworks based on gene expression data with a heuristic method to find top-scoring networks. Since then, multiple methods for solving this *ac-tivemoduleidentificationproblem* were developed. One of the most notable methods, called BioNet, was proposed by Dittrich *et al.* [4]. They suggested to use a maximum-likelihood-inspired subnetwork scoring scheme such that finding the best scoring subnetwork corresponds to solving the Maximum-Weight Connected Subgraph (MWCS) problem. While the problem is NP-hard, in the same paper a practical exact solver was proposed. Maximum-likelihood inspired formulation combined with an exact solver for the corresponding problem allowed to achieve great performance on both simulated and real data.

A question that is not usually addressed in the existing methods for solving the active module identification problem is the level of confidence of individual vertex inclusion. By design, in the resulting networks vertices with high individual significance are connected via less significant vertices. This raises the question whether vertices included in the module are more important compared to vertices with similar individual significance that are not included in the module. This is particularly important when an individually non-significant vertex is included in the module. Uncertainty in this aspect can lead to misinterpretation of the data either by attributing importance to false vertices or missing key vertices.

Previously Beisser et al [2] suggested a jackknife resampling approach where the active module problem is solved multiple times for resampled input data. It allows to introduce *support* values: how many times a particular vertex or edge was a part of the solution for the resampled data. The calculated support values can be then used to distinguish robust signals from noise in the resulting module. However, this method is limited to experiments with a large number of replicates and can not be applied for small-scale experiments.

The way different module confidence thresholds are handled is another problem of binary methods like BioNet. For example, a module of higher confidence could contain vertices not present in a module of lower confidence. Since in a real-case scenario several confidence thresholds are considered, such inconsistencies impede the interpretation of the results. To address this issue, previously [9] we considered a problem of connectivity-preserving vertex ranking – a ranking with the constraints that: 1) each prefix of the ranking should induce a connected subgraph, and 2) smaller induced subgraphs correspond to more confident modules. In that paper we proposed a semi-heuristic ranking method that was better (as measured by the area under the ROC curve, AUC ROC) compared to both baseline vertex ranking by individual input significance and ranking from multiple BioNet runs with different thresholds.

In the current study we consider the active module identification problem as a soft vertex classification problem. In this case, instead of providing a hard classification of vertices into either being in the module or not, we estimate probabilities of each vertex to belong to the active module. To estimate these probabilities we propose a method that is based on Markov chain Monte Carlo (MCMC) module sampling from a posteriori distribution. First, by producing the vertex probabilities, this approach directly resolves the question of individual vertices confidence. Moreover, we show that the estimated probabilities can be used to calculate expected AUC ROC of any ranking and thus the problem of finding the best connectivity-preserving ranking can be defined constructively. We prove that this problem is NP-complete, but show that in practice it can be solved accurately and efficiently using a heuristic algorithm. Finally, we show that the method can achieve high classification accuracy on simulated data and works well on real data.

## 2. Approach

While the active module problem appears in many different contexts, for the sake of clarity we focus here on the example of protein-protein interaction networks together with gene expression data. We formalize the protein-protein interaction network as a graph *G*, where vertices correspond to protein-encoding genes, and two vertices are linked by an edge if the corresponding proteins interact. Gene expression data are given for sets of samples for two biological conditions of interest (e.g. control and treatment), so that differential expression *p*-values can be calculated for each gene. While for some genes the null hypothesis of having zero expression change between conditions is true, and so the corresponding *p*-values are uniformly distributed on [0, 1], *p*-values for “interesting” genes that exhibit a difference in expression would tend to be closer to zero. According to [8], those interesting genes form a connected subgraph in *G*, which is called an active (or functional) module.

### 2.1. Beta-uniform mixture model

As shown in [14, 4], the distribution of all *p*-values can be approximated by the so-called beta-uniform mixture (BUM) distribution, where the beta component corresponds to a signal in the data, and the uniform component corresponds to noise. Let us recall that the beta distribution has support [0, 1] and is defined by its density

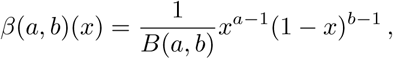

where *B*(*a,b*) is the Beta function. The BUM distribution is a mixture of uniform and beta *β*(*a,* 1) distributions and is defined by its density

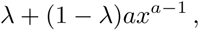

where *λ* is the mixture weight of the uniform component and *a* is the shape parameter of the beta distribution.

We assign a weight *w*(*v*) to each vertex *v* of the graph *G* equal to the *p*-value assigned to the corresponding gene. Thus, in our model we have: a connected graph *G* on *n* = *|G|* vertices, its connected subgraph *M*, a family of independent random variables *W*_*v*_, *v∈V* (*G*)\*V* (*M)* with uniform distribution on [0, 1], and a family of independent random variables *W*_*v*_, *v∈V* (*M)* with *β*(*a,* 1) distribution.

### 2.2. MCMC approach

Our goal is to find out which vertices are likely to belong to the active module*M* :

**Problem 1.** *GivenaconnectedgraphGandvertexweightsw*_*v*_ *∈* [0, 1] *findtheprobabilityP* (*v∈M|W* = *w*) *foreachvertexvtobelongtothemoduleM.*

We solve this problem in the following way. We generate a large sample S of random subgraphs *S* from conditional distribution *P* (*S* = *M|W* = *w*) using the Metropolis–Hastings algorithm (see Section 3). While all the probabilities *P* (*S* = *M|W* = *w*) are small and the multiplicity of each subgraph in S is small, the probabilities of each vertex to belong to the module are cumulative statistics and show robust behavior. The same holds for the probabilities *P* (*V⊂M|W* = *w*), where *V* are relatively small sets of vertices.

The benefits of the soft classification approach are:

1. it allows to estimate the level of confidence that a particular gene is expressed differently, which is the probability that a corresponding vertex belongs to the active module;
2. it allows to analyze alternative complement pathways by studying probabilities of the form *P* ((*V*_1_ *⊂M)V* (*V*_2_ *⊂M)|W* = *w*);
3. for any set of genes *V* reported as differently expressed, our method allows to compute the false discovery rate (FDR) as 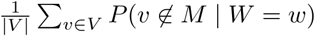;
4. it provides a vertex ranking that maximizes the area under the ROC curve (AUC ROC) (see Section 3.3);
5. it allows to heuristically find a vertex *connectivity preserving* ranking that maximizes the AUC ROC (see Section 3.4).

## 3. Methods

### 3.1. Markov chain Monte Carlo based method

We solve Problem 1 in the following way: we sample a set 𝕊 of random subgraphs *S* from conditional distribution *P* (*S* = *M|W* = *w*) using the Markov chain Monte Carlo (MCMC) approach, and estimate *P* (*v∈M|W* = *w*) as

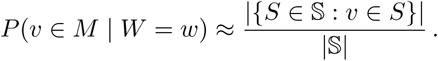

First, we estimate the beta-uniform mixture parameters 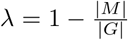 with a maximum-likelihood estimator [3].

For MCMC sampling we implement the Metropolis-Hastings algorithm [6]. It starts at a random subgraph *S*_0_ of order *k* = *|M|* = (1 *-λ*)*|G|*. On each step *i* we choose a candidate subgraph *S´* by removing a vertex *v*_*-*_ from *S*_*i*_ and adding another vertex *v*_+_ from the neighborhood of *S*_*i*_ (by neighborhood of a subgraph *S*

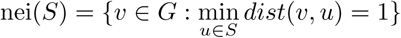

we mean the set of all vertices of *G* at distance 1 from *S*). We note that the subgraph *S´* always has the same number of vertices *k* as *S*_0_. The proposal probability *Q*(*S´|S*_*i*_) is

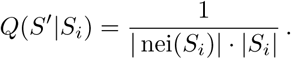

We set the acceptance probability in the Metropolis-Hastings algorithm as

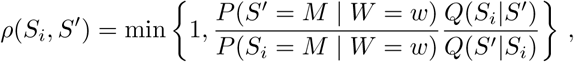

where *P* (*S´* = *MW* = *w*) = 0 as soon as *S´* is not connected.

For any connected subgraph *S* of *G* the value *P* (*S´* = *M|W* = *w*) can be expressed in terms of probability density *p* of an absolutely continuous random vector variable *W* :

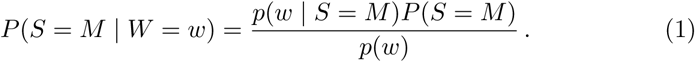

So,

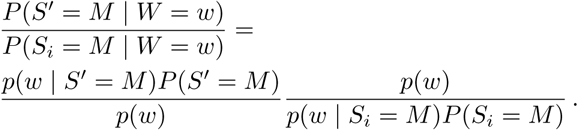

The fraction 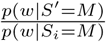 is equal to

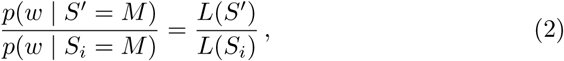

where *L*(*S*) is the likelihood of the subgraph *S*. In our case

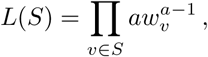

so, we have:

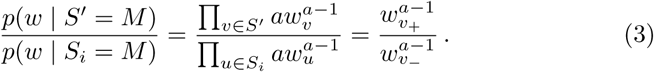

Assuming that the prior distribution of choosing the module *P* (*S* = *M)* is uniform^1^ on the set of connected subgraphs *S* of the same order *|M|*, we get

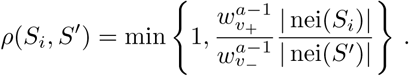

The proposed MCMC algorithm can visit all possible subgraphs *S*, and so it converges to the distribution *P* (*S* = *M|W* = *w*).

**Algorithm 1:**
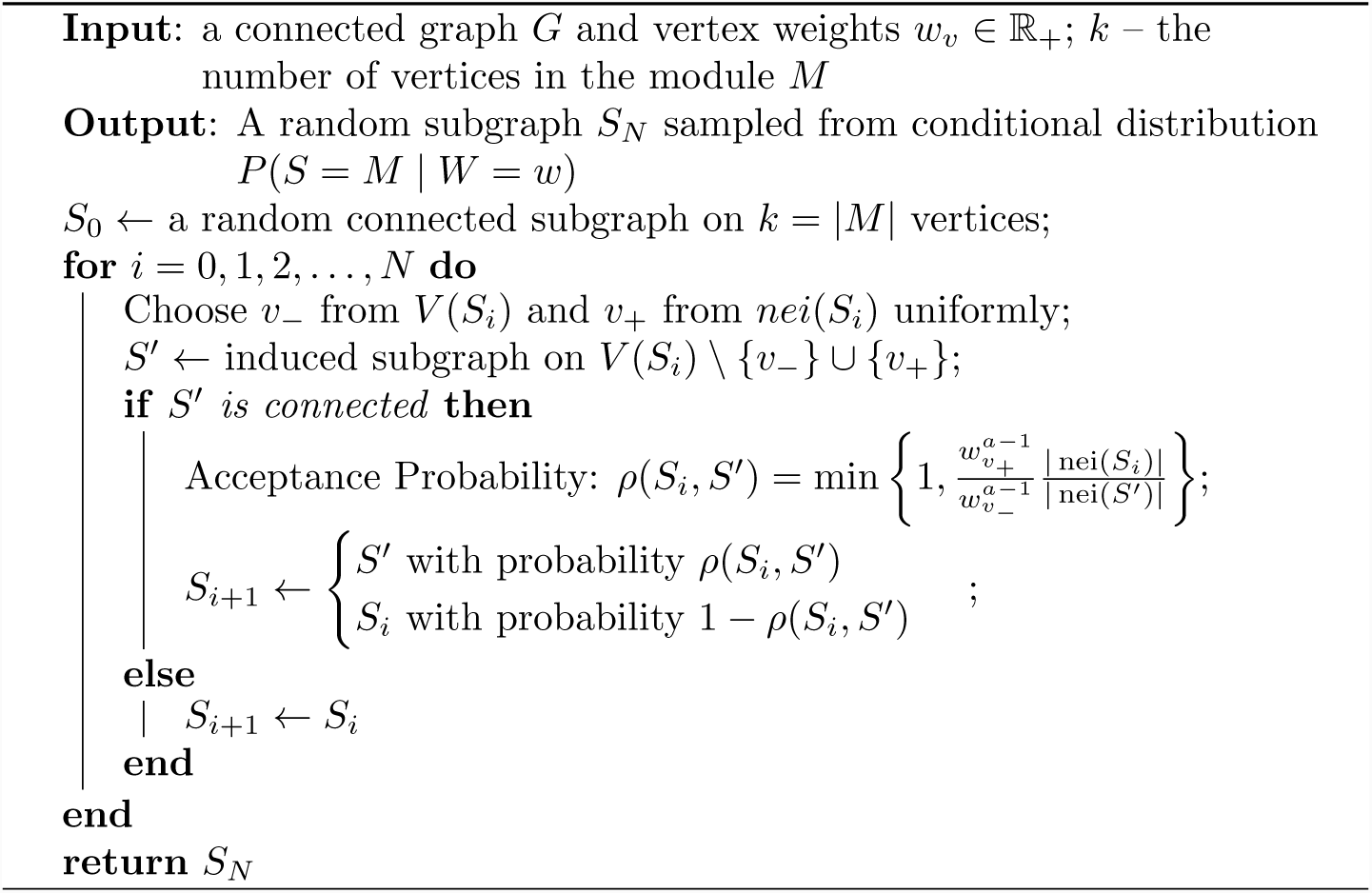
Metropolis-Hastings algorithm

### 3.2 Heuristic approach to arbitrary module order

Since on the real data the estimated order of an active module has a tendency to be too large (1 − *λ* is up to 0.5), we adjust our method in order to allow to change the number of vertices in the subgraph during the MCMC process. In this approach on each step of the process one can either add one vertex to the subgraph or remove one vertex from it. In order to provide subgraphs of biologically relevant size, we penalize each additional vertex by a factor *aτ* ^*a-*1^, where *τ* is some confidence threshold as described in [4]. More formally, it means that we consider such a prior distribution on subgraphs that 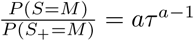 where *S*_+_ is an induced subgraph on the vertices *V* (*S*) *∪*{*v*_+_}. Thus our heuristic approach is very similar to Algorithm 1, but

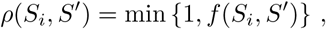

Where

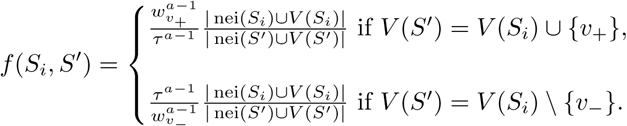

**Algorithm 2:**
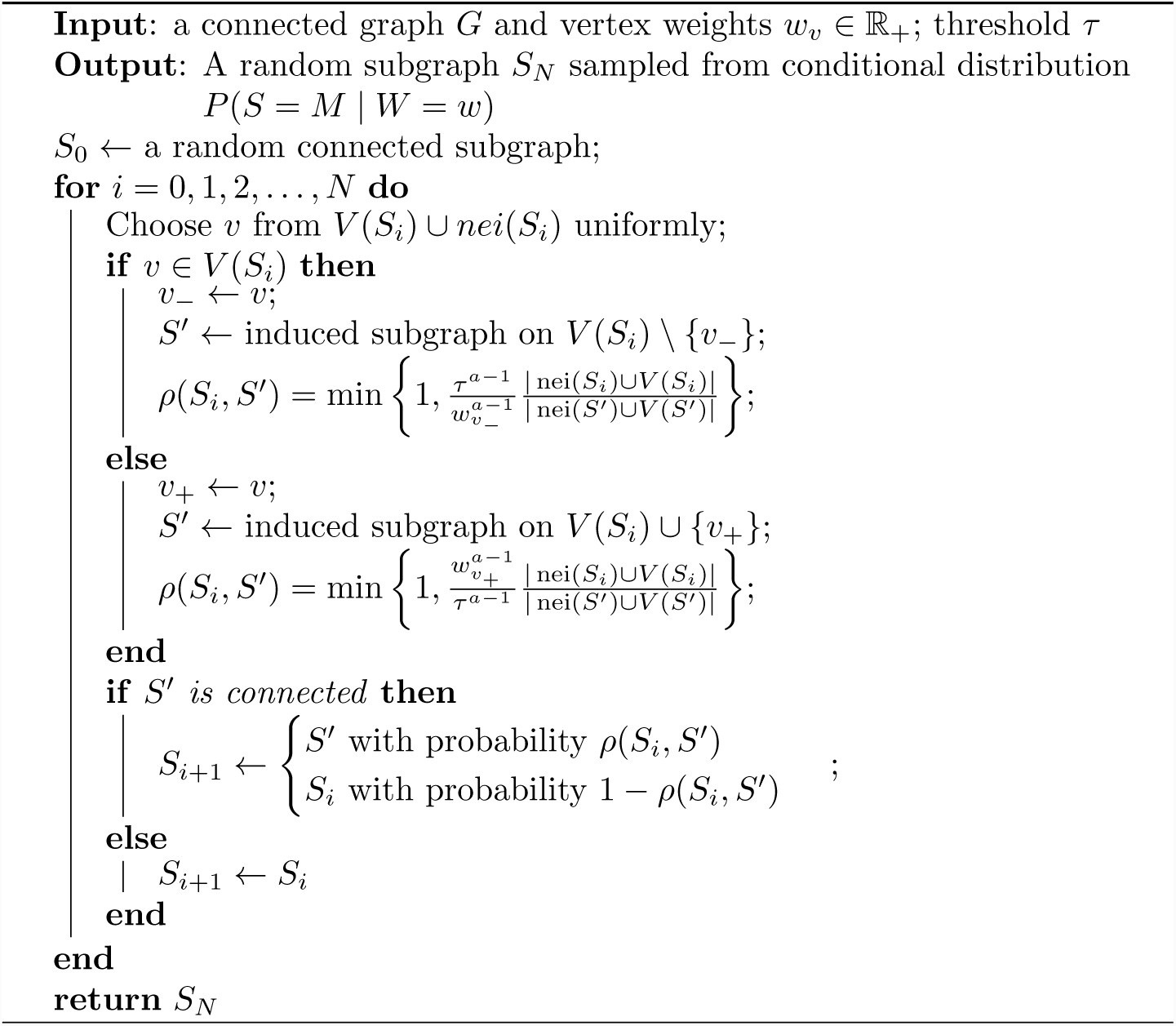
Metropolis-Hastings algorithm for subgraphs with an arbitrary number of vertices

For estimating probabilities *P* (*v∈M|W* = *w*) we need to choose a set of subgraph samples 𝕊. Here we consider two ways of doing this. For both ways we need to estimate the mixing time *T* of the algorithm: the number of Markov chain iterations such that the distribution of *S*_*T*_ approximates the target distribution well. We note that theoretically the Markov chain converges to the desired distribution since it is ergodic: it is recurrent (since the number of states is finite), it is aperiodic (since there is a positive probability that the process stays at the same state), and it is irreducible since by construction it can reach any subgraph from any other subgraph. In practice, the mixing time depends on multiple parameters including the graph *G* order. The first way to choose S is to do a number of independent MCMC runs of *T* iterations and add each *S*_*T*_ to S. Here, all samples in S are independent, which can be used to calculate the accuracy of the vertex probabilities estimation. Another way consists in doing one long run of MCMC and putting all *S*_*i*_ for *i>T* into S. Although consecutive samples are not independent, the probabilities that are estimated this way converge to the true probabilities given sufficiently long series (see Section 4.1 for discussion of the converging time on the simulated data).

### 3.3 Features of soft classification solution

Having the set S of subgraphs sampled from the conditional distribution *P* (*S* = *M|W* = *w*), one can easily answer the following questions. First, one can estimate for each vertex *v*_*i*_ the probability *p*_*i*_ that it belongs to the active module. In the case when the module size is not fixed, we can also estimate the conditional probabilities

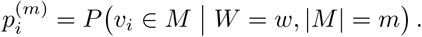

Second, for any boolean-valued function *A*(*M)* (for example, *A*(*M)* = (*v*_1_ *∈M)* (*v*_2_ *M)∧* ¬ (*v*_3_ ∈ *M)*) one can estimate the probability *P* (*A*(*M)|W* = *w*).

Note, that if each *S* from 𝕊 was generated with an independent MCMC run, then each *A*(*S*) can be considered to be a result of a Bernoulli trial. Thus, all such probabilities can be estimated with a mean square error of the order 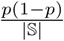, where *p* = *P* (*A*(*M)|W* = *w*).

We also show how the probabilities of each vertex to belong to the active module are related to the expected AUC ROC for some vertex ranking. For a vertex ranking *v*_1_, *v*_2_,*…,v*_*n*_, where *n* = *|G|*, and a module *M* of order *m* the ROC curve is a step-curve in a square [0, 1] *×* [0, 1], which starts at(0, 0), and on the *i*-th step it goes either up 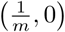 (if *v*_*i*_ *∈M*) or right 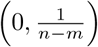 The AUC ROC shows how accurate the classification is.

We prove the following lemma:

#### Lemma 1.

*Let n be the number of vertices in G, m be the number of vertices in the module M, v*_1_, *v*_2_,*…*, *v*_*n*_ *be some vertex ranking and p*_*i*_ *be the probabilities p*_*i*_ = *P* (*v*_*i*_ *∈M|W* = *w*). *Then the expected value of the AUC ROC is equal to*

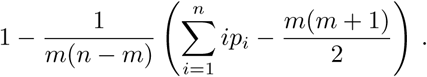

*Proof.* See Section S1 in Supplementary Materials.

Lemma 3 implies that the vertex ranking with the largest expected value of the AUC ROC is the ranking according to a descending order on *p*_*i*_ (for any particular module order *m*). If the number *m* of the vertices in the module is not known, the expected AUC ROC is equal to

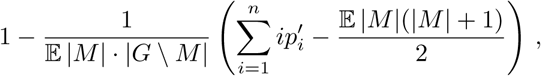

Where

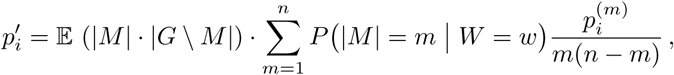

and 𝔼 stands for the expected value.

We note that the probabilities 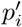 can be estimated based on the sample 𝕊 or approximated by *p*_*i*_.

### 3.4 Connectivity preserving ranking

While we provide the best solution for the soft classification problem in terms of the expected AUC ROC, one can be interested in getting the solution for the hard classification problem as well. Our method is easily transformable into a hard classification method. Namely, our goal is to define a module *M* (*q*) for any FDR level *q* in a consistent manner. That is, we don’t allow the situation when a vertex is included in a module with a small FDR level, but excluded from a module with a larger FDR (formally speaking, we demand that *M* (*q*_1_) *⊂M* (*q*_2_) as soon as *q*_1_ *<q*_2_). Any such family of modules *M* (*q*) corresponds to a connectivity preserving ranking.

**Definition 1.** *For a graph G with vertices v*_*i*_ *the* connectivity preserving *rank-ing is such a ranking of v*_*i*_ *that for any k-prefix v*_1_, *v*_2_,*…,v*_*k*_ *the induced graph on this set of vertices is connected.*

We want to choose the best connectivity preserving ranking in terms of the expected value of the AUC ROC. Since the expected AUC ROC depends only on *p*_*i*_ (or their modified version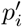), we can formulate the following problem:

**Problem 2** (Optimal Connectivity Preserving Ranking, OCPR). *GivenagraphGandthelistofitsverticesv*_*i*_, *eachequippedwiththeprobabilityp*_*i*_, *findthe* connectivity preserving *rankingmaximizingtheexpectedAUCROC.*

#### Lemma 2.

*OCPRproblemisNP-hard.*

*Proof.* See Section S2 in Supplementary Materials.

As Problem 2 is NP-hard, here we provide a heuristic algorithm to solve it. On the *k*-th step of the algorithm we have an integer number *r*_*k*_ and *G*_*k*_ – a connected subgraph of *G* (where *r*_1_ = *n* and *G*_1_ = *G*), and we define a connected subgraph *G*_*k*+1_ in the following way. For each vertex *v* we define a *k*+1 *k*+1 subgraph 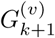 as the largest connected component of *G*_*k*_ *\*{*v*} and 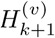 as 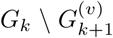. Then we choose such a vertex *v* that maximizes average FDR in 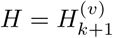, which is equal to 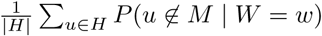. Then we assign to all vertices in the subgraph *H* the rank *r*_*k*_ and assign to *r*_*k*+1_ = *r*_*k*_ *|H|*

Each step requires time *O*(*n*^2^), so the performance time is *O*(*n*^3^). On our real data example, the running time was 13 seconds on Intel(R) Core(TM) i5-7200U CPU @ 2.50GHz (implemented in C++).

### 3.5 Baseline ranking methods

We compared our approach with three other ranking methods on simulated data in Section 4.1. The first method ranks vertices by the ascending order of their input weights *w*_*v*_. In the second method vertices are ranked by the descending order of the number of occurrences in the modules as found by running the BioNet method with 20 different significance thresholds. The thresholds are selected to be distributed at equal intervals between maximum and minimum vertex log-likelihoods. The third method is a semi-heuristic ranking method from [9] developed for finding a good connectivity preserving ranking. To describe it briefly, it first finds the maximum-likelihood connected subgraph and recursively ranks a set of vertices inside and outside of the subgraph. The recursion step also involves finding a maximum-likelihood connected subgraph but with constraint on consistent connectivity with already defined ranking.

Note that we do not compare our method with the jackknife resampling method [2] on simulated data as it requires simulating gene expression data, which can induce artificial biases in the evaluation. Instead, we compare our results with the results of this method on real data (see Section 4.2).

## 4. Discussion

### 4.1. Experiments on simulated data

First, we consider a toy example. We generated a random graph *G* on 30 vertices and 65 edges. Then we chose the active module *M* on 9 vertices randomly and generated the weights from the beta distribution *β*(0.2, 1) for the vertices in *M* and from the uniform distribution for all other vertices. For such a toy example we can both compute the probabilities *P* (*v∈M|W* = *w*) directly from (1) and estimate them as 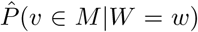 using our MCMC approach. We run 10^6^ iterations of the Metropolis-Hastings algorithm. The estimations approximated the actual probabilities very accurately with root-mean-square error equal to 2 · 10^*-*3^ We also note that the ranking of vertices based on estimations 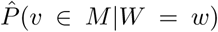 was the same as the ranking based on true probabilities. For both actual and estimated probabilities the AUC ROC is equal to 0.92.

Second, we compared our MCMC-based connectivity preserving ranking method with three other ranking methods (see Section 3.5) on simulated data. Here we considered 50 random instances. For each instance a random scale-free graph on 100 vertices was generated. Then we generated active modules in two steps: 1) the number of the vertices in a module was uniformly selected from 5 to 25 and 2) a module was chosen uniformly at random from all connected subgraphs with the selected order. Vertex weights were generated from the beta-uniform mixture distribution with values of the *a* parameter selected from the [0.01, 0.5] interval. The AUC measures for rankings produced by four tested methods are shown on Fig. 2. For all values of *a* our approach shows significantly better performance compared to all three baseline methods.

Next we considered MCMC performance on a real-world protein-protein interaction graph. We used a graph with 2,034 vertices and 8,399 edges as constructed in [4] for a diffuse large B-cell lymphoma dataset. We chose an active module uniformly at random from connected subgraphs on 200 vertices. Weights of the vertices in the active module were generated from the beta distribution *β*(0.25, 1).

First, we considered the behavior of log-likelihood values for samples during one MCMC run (Fig. 1, green line). This plot shows that the log-likelihood value stabilizes after about 25,000 iterations. Thus, we can estimate the MCMC mixing time *T* as 25,000 for this case. Then we checked that 25,000 iterations is sufficient for estimating vertex probabilities. For different values of *T´* we calculated AUC values for rankings based on 1,000 independent runs of MCMC for *T´* iterations (Fig. 1, red line). The results show that indeed 25,000 iterations is enough to achieve high AUC values, and the saturation phase begins even earlier. Finally, we compared ranking for one long MCMC run and for 1,000 independent samples (Fig. 1, blue line). For one long run we estimated probabilities using all generated MCMC samples except the first 25,000. It can be seen that AUC values saturate after about 50,000 total MCMC iterations which justifies the usage of one long MCMC run. Practically, this means that good probability estimates can be achieved very fast, given that 100,000 MCMC iterations took about one minute on a laptop.

**Figure 1:**
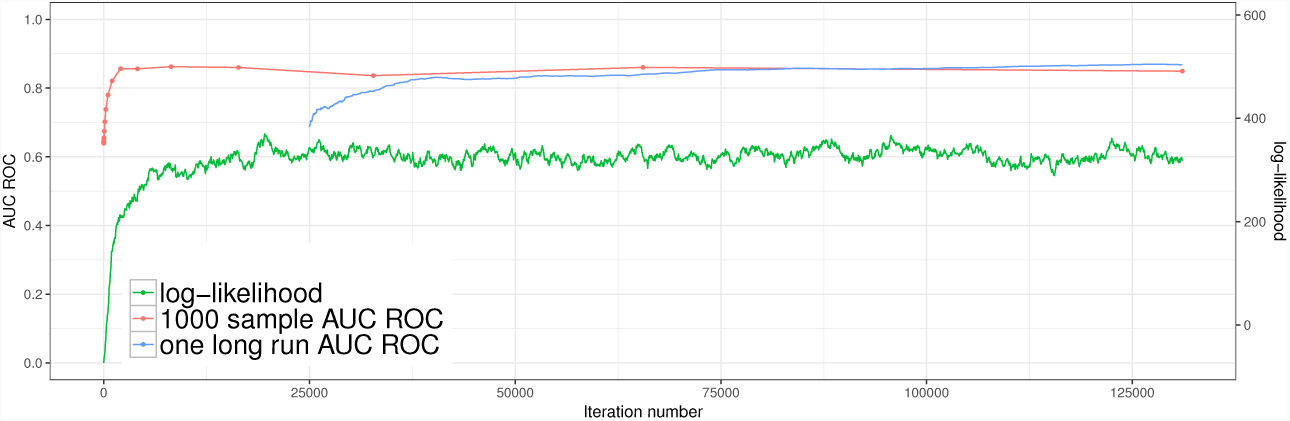
Behavior of subgraph log-likelihood values and ranking AUCs depending on the number of MCMC iterations. Real protein-protein interaction graph of 2,034 vertices is used as *G*, module of 200 vertices is chosen uniformly at random. Green line: log-likelihood values for subgraphs *S*_*i*_ generated during one MCMC run. Red line: AUC values for rankings based on 1,000 independent MCMC samples depending on the chosen mixing time estimate. Blue line: AUC values for rankings based on one MCMC run calculated on all samples *S*_*i*_ for *i >* 25,000.

**Figure 2:**
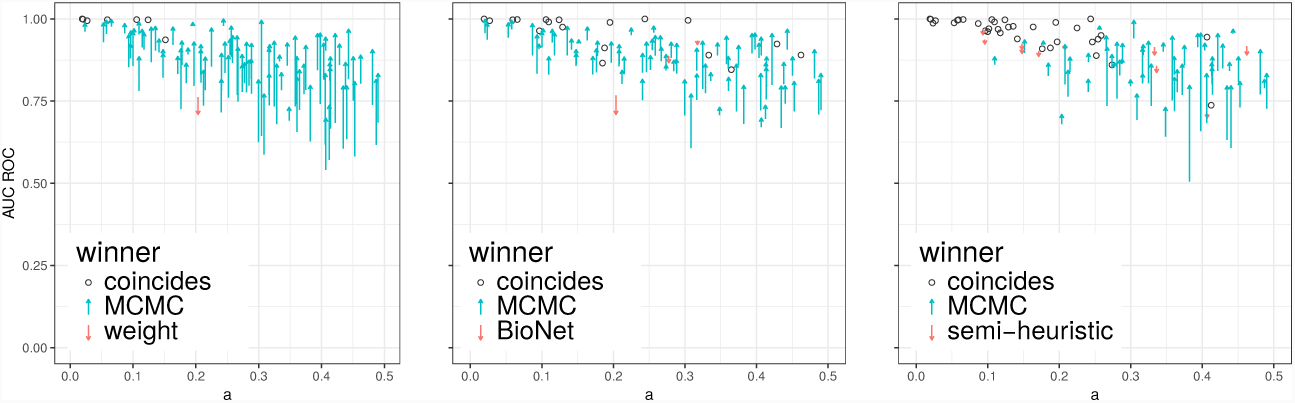
Ranking AUC values for simulated instances for graphs on 100 vertices. The proposed MCMC method is compared with the following methods: ranking by input weights (left), BioNet-based ranking (middle) and semi-heuristic ranking from [9] (right). One arrow corresponds to one experiment. The head of each arrow points to the AUC score of the MCMC method, while the tail indicates the AUC score of the corresponding alternative method. Thus, upward arrows indicate instances in which the MCMC method has a higher AUC score. Dots indicate the instances for which the results are equal. Color depends on which method has better AUC.

### 4.2 Experimental results on real data

We apply our method to the diffuse large B-cell lymphoma dataset and the protein-protein interaction graph constructed in [4]. The *p*-values in the DLBCL dataset are the result of a differential expression *t*-test between the two tumor subgroups: germinal center B-cell-like (GCB) DLBCL and activated B-cell-like (ABC) DLBCL. The estimated BUM distribution parameters are *λ* = 0.48 and *a* = 0.18. As described in Section 3.2, we penalized the addition of a new vertex to the module using the confidence threshold *τ* = 10^*-*7^.

The obtained module is shown on Fig. 3. The module shows the prefix of the ranking with FDR = 0.25. Additionally, we highlighted submodules of more strict FDR values: 0.15 and 0.05. Note that, by construction, a more strict module is a subgraph of a less strict one. The genes in the modules are mostly known to be associated with cancer. For example, genes BCL2, BMF, CASP3, PTK2 and WEE1 are involved in the cell apoptosis pathway. Some other genes, like LYN or KCNA3, are associated with proliferation and signalling in hematopoietic cells.

**Figure 3:**
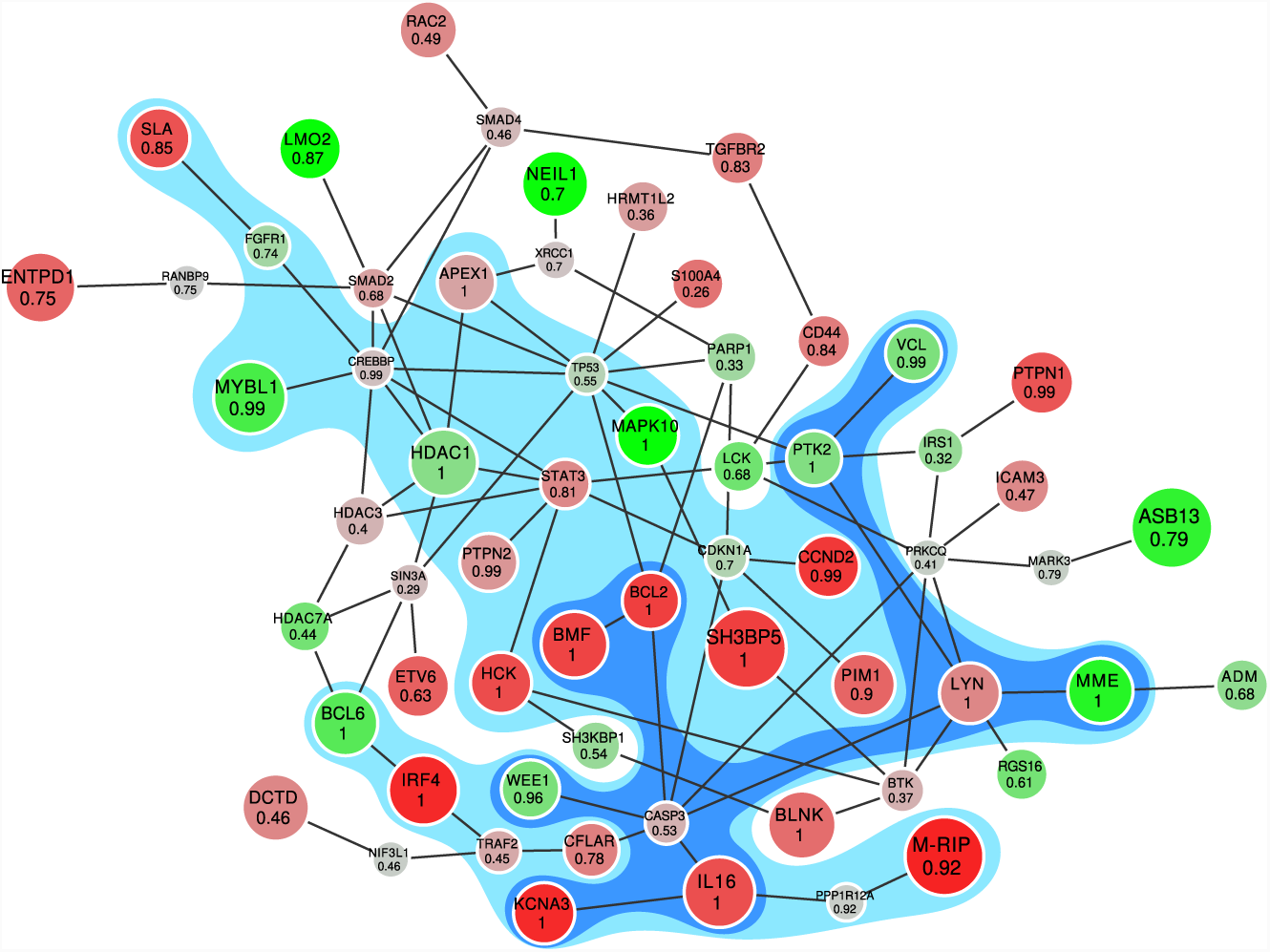
The module for comparison of GCB and ABC types of diffuse large B-cell lymphoma (red vertices are up-regulated in ABC type, green ones are up-regulated in GCB). The vertices of the blue subgraph belong to the active module with very high confidence (FDR is 0.05). The vertices of the light blue subgraph belong to the active module very likely (FDR is 0.15). The number in each vertex means the frequency of the vertex presence in a sampled subgraph.

Additionally, we compared our result to one obtained by the jackknife resampling procedure as described in [2] with 100 resamples and the same threshold of *τ* = 10^*-*7^. Resampling support values are consistent with MCMC-based probabilities (see Supplementary material). However, compared to the resampling method [2] our approach has two major advantages. First, out method starts with *p*-values and can be used even for experiments with a small number of replicates, where resampling is not feasible. Second, calculating individual vertex probabilities does not involve solving NP-hard problems and can be easily done in practice without dependency on external solver libraries like IBM ILOG CPLEX. Even the NP-hard problem of finding the AUC ROC optimal connectivity preserving ranking can be solved well on real data with the heuristic algorithm.

## 5. Conclusion

While the active module identification problem was intensively studied in recent years, most of the approaches solve it as a hard classification problem. Here we study the question of assigning confidence values for individual vertices and thus consider a soft classification problem. All the hard classification methods could only provide a blackbox which tells us whether a particular gene belongs to the active module or not, and so, by design, they all have a flaw: they don’t provide any way to distinguish between more and less confident inclusions. The soft classification approach addresses this issue.

We propose a method to estimate probabilities of each vertex to belong to the active module based on Markov chain Monte Carlo sampling. Based on these probabilities, for any given FDR level we can provide a solution to the hard classification problem in a consistent manner: a module for a more strict FDR level is a subgraph of any module for a more relaxed FDR level. Overall, the proposed approach is very flexible: it starts with *p*-values, and thus can be used in many different contexts, and does not depend on external libraries for solving NP-hard problems.

## Acknowledgements

The authors thank Artem Vasilyev for fruitful discussions.

## Supplementary Materials S1

In this section we prove Lemma 3.

### Lemma 3.

*Let n be the number of vertices in G, m be the number of vertices in the module M, v*_1_, *v*_2_,*…*, *v*_*n*_ *be some vertex ranking and p*_*i*_ *be the probabilitie sp*_*i*_ = *P* (*v*_*i*_ *∈M|W* = *w*). *Then the expected value of the AUC ROC is equal to*

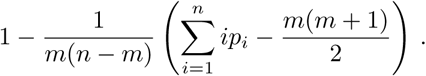

*Proof.* First of all, we introduce a rotated not-normalized ROC curve (Fig.4). On the *i*-th step this curve goes either up 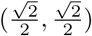 (if *v*_*i*_ *∈M)* or down 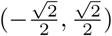 (if *v*_*i*_ *∉M)*

This linear transformation of the ROC curve is a function, so one can find not only the expected value of the area under it, but its pointwise expected value as well. Let its height at the point 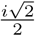 be *h*_*i*_. Then the expected value of *h*_*i*+1_ *-h*_*i*_ is 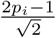. The area under the (original) ROC curve (up to the factor *m*(*n−m*)) is equal to the area under the rotated ROC curve plus the area of the quadrilateral *AECD*. The area of *AECD* is equal to the area of *ABCD* (which is equal to *m*(*n-m*)) minus the area of *ABF* (which is equal to 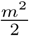) plus the area of *CEF* (which is equal to 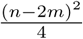. Thus, we obtian:

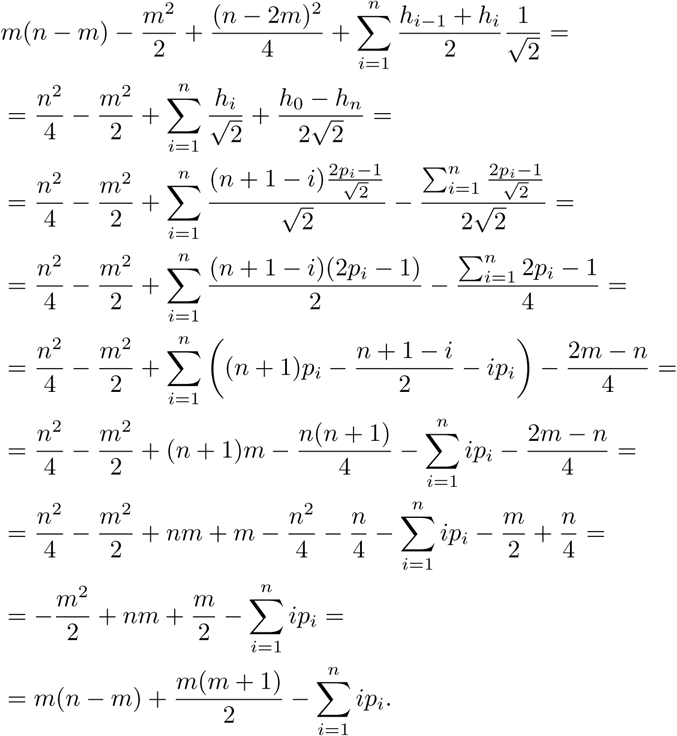

After normalization we obtain

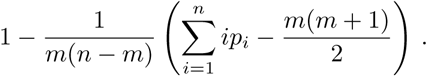

## S2

In this section we prove Lemma 4.

### Lemma 4.

*TheOCPRproblemisNP-hard.*

*Proof.* To prove that OCPR is NP-complete, we will use a modification of the method for proving that the Steiner tree problem in graphs is NP-complete. The referenced method is described in [16].

In decisional form, OCPR is equivalent to:

OCPR (decisional). *Given a graph G, the list of its vertices v*_*i*_, *each equipped with the probability p*_*i*_, *and a real numberk, determine if there is a* connectivity preserving *rankingv*_1_, *v*_2_,*…*, *v*_*n*_ *such that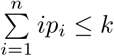*

First, we need to show that OCPR is in NP. Given an hypothetic positive solution *v*_1_, *v*_2_,*…*, *v*_*n*_, it is trivial to check in polynomial time that this ranking is *connectivity preserving* and 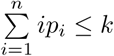 holds.

The Exact Cover by 3-Sets problem (X3C) is a well-known NP-complete problem mentioned in [5]:

Problem X3C. *Given a finite set X with |X|* = 3*q and a collection C of 3-element subsets of X (C* = {*C*_1_,*…*, *C*_*n*_}, *C*_*i*_⊆ *X,|C*_*i*_*|* = 3 *for* 1 *in), determine if C contains an* exact cover *for X, that is, a subcollection C*´ ⊆ *such that every element of X occurs in exactly one member of C* ´*?*

Let us propose a reduction from X3C to OCPR giving a set of rules to build an instance of OCPR starting from a generic instance of X3C. If we prove that this transformation is executable in polynomial time, we will also prove that OCPR is NP-complete.

Given an instance of X3C, defined by the set *X* = {*x*_1_,*…,x*_3 sub>*q*_} and a collection of 3-element sets *C* = { _1_,*…,C*_*n*_}, we have to build an OCPR instance specifying the graph *G* = (*V,E*), the probabilities *p*_*i*_, and the upper bound on the required sum *k*.

The vertices of *G* are defined as:

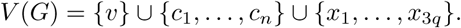

That is, we add a new node *v*, a node for each member of *C*, and a node for each element of *X*.

The edges of *G* are defined as:

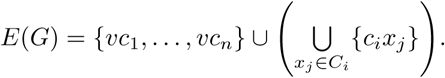

That is, there is an edge from *v* to each node *c*_*i*_, and an edge *c*_*i*_*x*_*j*_ if the element *x*_*j*_ belongs to the set *C*_*i*_ of the X3C instance.

Let 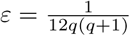 The probabilities *p*_*i*_ are defined as:

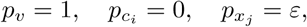

and the upper bound on the required sum is defined as 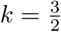.

The reduction from X3C to OCPR is easy to do in polynomial time.

**Proposition 1.** *There exists a connectivity preserving ranking with 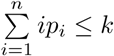 if and only if there is an exact cover in the corresponding instance of X3C.*

**Proof.** We split the proof into two parts, one for each implication.

- X3C *⇒* OCPR Suppose there is an exact cover *C´* for the X3C problem. Clearly, *C*^*t*^ uses exactly *q* subsets. Without loss of generality suppose they are *C*_1_,*…*, *C*_*q*_ (if it is not the case, we just have to relabel them). Then a valid connectivity preserving ranking looks as follows:

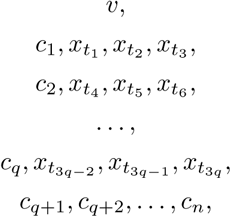

where every *C*_*i*_ consists of elements 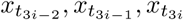. Then:

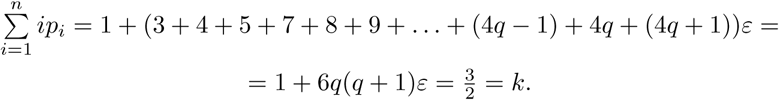
- X3C *⇐* OCPR Suppose there is a connectivity preserving ranking with 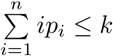 Since *k<* 2 and *p*_*v*_ = 1, the first vertex in this ranking must be *v*. Let 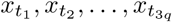 be the order the vertices corresponding to the X3C instance elements appear in this ranking. For every *j*, vertex *v* and at least 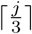 vertices corresponding to 3-element sets of the X3C instance must appear before vertex 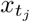 in the ranking, since every vertex *c*_*i*_ appearing in the ranking opens the way for at most three new vertices *x*_*j*_ to appear afterwards. Hence, the earliest position vertex 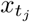 might appear at in the ranking is 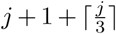. But if every 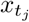 does appear at its earliest possible position, then, like in the former case,

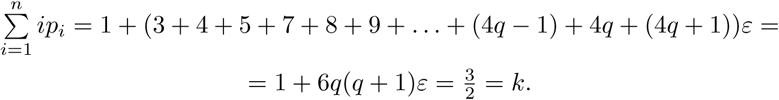 If any vertex 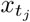 appears at a later position, its index in the ranking will increase, the sum of *ip*_*i*_ will increase and exceed *k* (since the sum of *ip*_*i*_ is already equal to *k* and 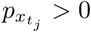). Therefore, every vertex 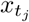 must appear at position 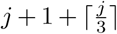, and the ranking looks like in the former case, too:

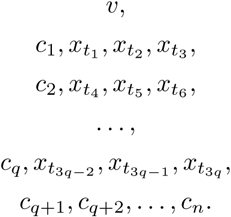 It’s easy to prove that 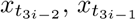 and 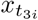 belong to *C*_*i*_ using induction on *i* from 1 to *q*. Hence, *C*^*t*^ = *C*_1_, *C*_2_,*…*, *C*_*q*_ is an exact cover for the X3C instance.

This concludes the proof.

## S3

In this section we compare our method and the jackknife resampling method described in [3]. We apply our method and the jackknife resampling method to the diffuse large B-cell lymphoma dataset and the protein-protein interaction graph constructed in [4]. The *p*-values in the DLBCL dataset are the result of a differential expression *t*-test between the two tumor subgroups: germinal center B-cell-like (GCB) DLBCL and activated B-cell-like (ABC) DLBCL. In our method we penalized adding a new vertex to the module using the confidence threshold *τ* = 10^*-*7^. For the jackknife method the *p*-values are recalculated for each of 100 data resamples and then for each collection of the *p*-values the active module identification problem is solved with BioNet (with the same threshold *τ* = 10^*-*7^). As the result of this procedure, for each vertex the support value (the occurrence frequency of the vertex in the solution) is computed. As one can see (Fig. 5), the results of these two methods are consistent.

**Figure 4:**
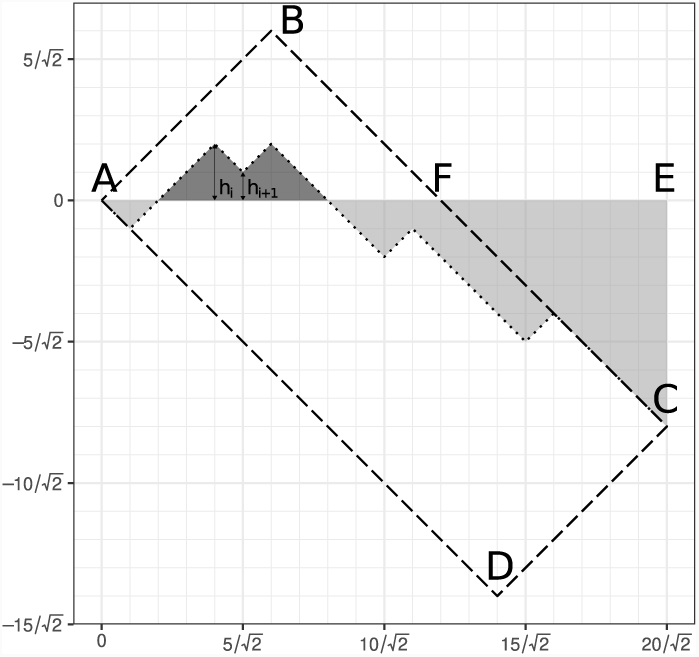
Rotated ROC curve

**Figure 5:**
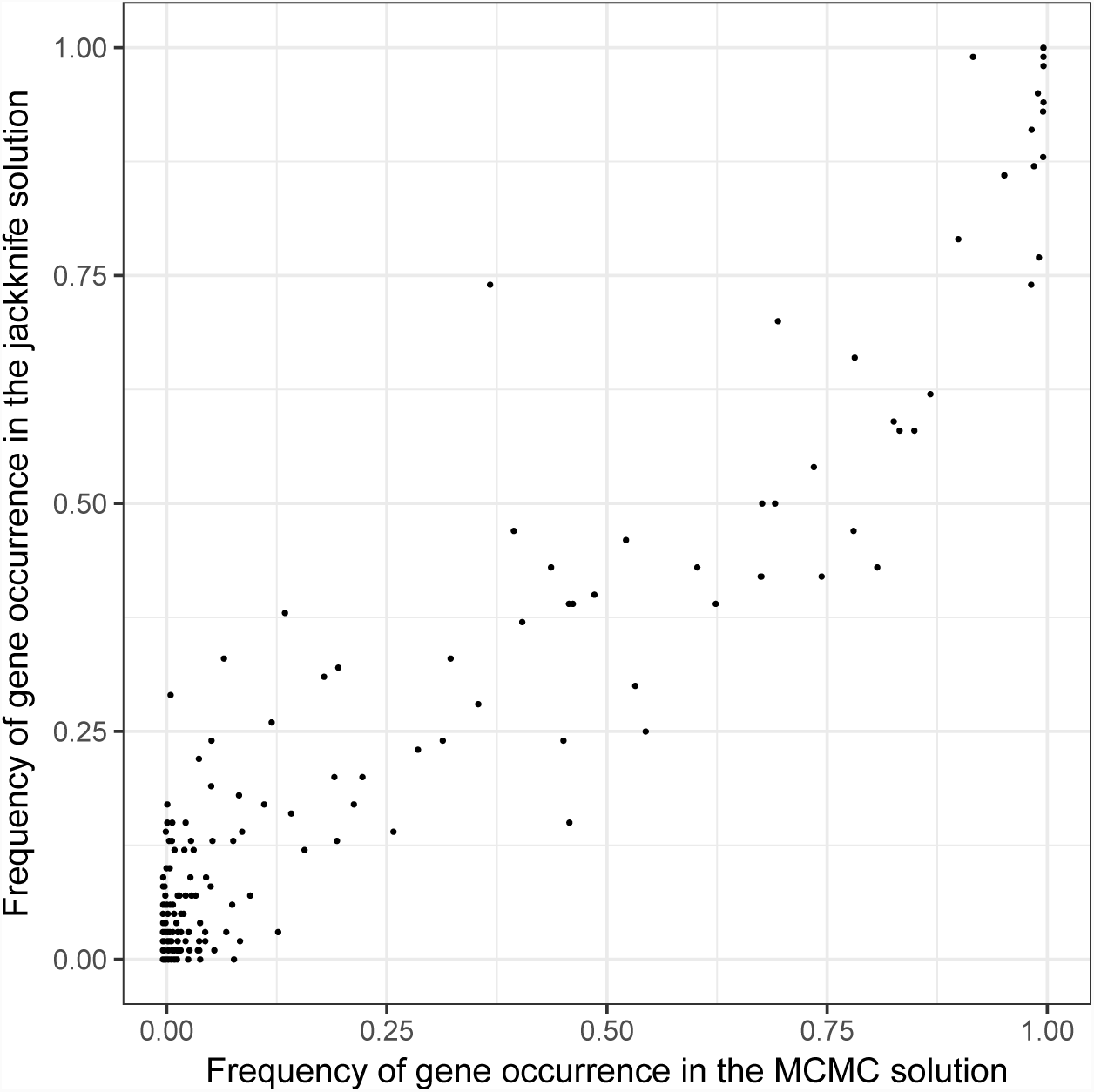
The support values and the probabilities of each vertex to belong to the active module are highly correlated.

## Footnotes

1 Since we do not make any additional assumptions about the module, the uniform distribution on subgraphs of equal order is the best choice of the prior distribution.

## References

[1] A. Alexeyenko, W. Lee, M. Pernemalm, J. Guegan, P. Dessen, V. Lazar, J. Lehtio, and Y. Pawitan. Network enrichment analysis: extension of gene-set enrichment analysis to gene networks. BMC Bioinformatics, 13:226, Sep 2012.

[2] D. Beisser, S. Brunkhorst, T. Dandekar, G. W. Klau, M. T. Dittrich, and T. Muller. Robustness and accuracy of functional modules in integrated network analysis. Bioinformatics, 28(14):1887–1894, Jul 2012.

[3] D. Beisser, G. W. Klau, T. Dandekar, T. Muller, and M. T. Dittrich. BioNet: an R-Package for the functional analysis of biological networks. Bioinformatics, 26(8):1129–1130, Apr 2010.

[4] Marcus T Dittrich, Gunnar W Klau, Andreas Rosenwald, Thomas Dandekar, and Tobias Müller. Identifying functional modules in protein-protein interaction networks: an integrated exact approach. Bioinformatics (Oxford, England), 24(13):i223–31, 2008.

[5] Michael R. Garey and David S. Johnson. Computers and Intractability: A Guide to the Theory of NP-Completeness. W.H. Freeman and Company, New York, 1979.

[6] W Keith Hastings. Monte Carlo sampling methods using Markov chains and their applications. Biometrika, 57(1):97–109, 1970.

[7] T. Ideker and N. J. Krogan. Differential network biology. Mol. Syst. Biol., 8:565, Jan 2012.

[8] Trey Ideker, Owen Ozier, Benno Schwikowski, and Andrew F Siegel. Discovering regulatory and signalling circuits in molecular interaction networks. Bioinformatics (Oxford, England), 18 Suppl 1:S233–S240, 2002.

[9] Javlon E Isomurodov, Alexander A Loboda, and Alexey A Sergushichev. Ranking vertices for active module recovery problem. In International Conference on Algorithms for Computational Biology, pp. 75–84. Springer, 2017.

[10] A. K. Jha, S. C. Huang, A. Sergushichev, V. Lampropoulou, Y. Ivanova, E. Loginicheva, K. Chmielewski, K. M. Stewart, J. Ashall, B. Everts, E. J. Pearce, E. M. Driggers, and M. N. Artyomov. Network integration of parallel metabolic and transcriptional data reveals metabolic modules that regulate macrophage polarization. Immunity, 42(3):419–430, 2015.

[11] A. Karnovsky, T. Weymouth, T. Hull, V. G. Tarcea, G. Scardoni, C. Laudanna, M. A. Sartor, K. A. Stringer, H. V. Jagadish, C. Burant, B. Athey, and G. S. Omenn. Metscape 2 bioinformatics tool for the analysis and visualization of metabolomics and gene expression data. Bioinformatics, 28(3):373–380, Feb 2012.

[12] M.D. Leiserson and others. Pan-cancer network analysis identifies combinations of rare somatic mutations across pathways and protein complexes. Nat. Genet., 47(2):106–114, Feb 2015.

[13] K. Mitra, A. R. Carvunis, S. K. Ramesh, and T. Ideker. Integrative approaches for finding modular structure in biological networks. Nat. Rev. Genet., 14(10):719–732, Oct 2013.

[14] S. Pounds and S. W. Morris. Estimating the occurrence of false positives and false negatives in microarray studies by approximating and partitioning the empirical distribution of p-values. Bioinformatics, 19(10):1236–1242, Jul 2003.

[15] E. J. Rossin and others. Proteins encoded in genomic regions associated with immune-mediated disease physically interact and suggest underlying biology. PLoS Genet., 7(1):e1001273, Jan 2011.

[16] Alessandro Santuari. Steiner tree np-completeness proof. Technical report, University of Trento, 2003.

